# Comparison of CRISPR/Cas endonucleases for *in vivo* retinal gene editing

**DOI:** 10.1101/2020.06.09.141705

**Authors:** Fan Li, Kristof Wing, Jiang-Hui Wang, Chi D. Luu, James A. Bender, Jinying Chen, Qi Wang, Qinyi Lu, Minh Thuan Nguyen Tran, Kaylene M Young, Raymond C.B. Wong, Alice Pébay, Anthony L. Cook, Sandy S.C. Hung, Guei-Sheung Liu, Alex W. Hewitt

## Abstract

CRISPR/Cas has opened the prospect of direct gene correction therapy for some inherited retinal diseases. Previous work has demonstrated the utility of adeno-associated virus (AAV) mediated delivery to retinal cells *in vivo*; however, with the expanding repertoire of CRISPR/Cas endonucleases, it is not clear which of these are most efficacious for retinal editing *in vivo*. We sought to compare CRISPR/Cas endonuclease activity using both single and dual AAV delivery strategies for gene editing in retinal cells. Plasmids of a dual vector system with SpCas9, SaCas9, Cas12a, CjCas9 and sgRNA targeting *YFP* and a single vector system with SaCas9/YFP sgRNA were generated and validated in YFP-expressing HEK293A cell by flow cytometry and T7E1 assay. Paired CRISPR/Cas endonuclease and its best performing sgRNA was then packaged into an AAV2 capsid derivative, AAV7m8, and injected intravitreally into CMV-Cre::Rosa26-YFP mice. SpCas9 and Cas12a achieved better knockout efficiency than SaCas9 and CjCas9. Moreover, no significant difference in *YFP* gene editing was found between single and dual CRISPR/SaCas9 vector systems. With a marked reduction of YFP-positive retinal cells, AAV7m8 delivered SpCas9 was found to have the highest knockout efficacy among all investigated endonucleases. We demonstrate that the AAV7m8-mediated delivery of CRISPR/SpCas9 construct achieves the most efficient gene modification in neurosensory retinal cells *in vitro* and *in vivo*.

## INTRODUCTION

Being discovered as a critical component of some bacterial and archaea, acting to counter viral intrusion (Jinek et al., 2012), the Clustered Regularly Interspaced Short Palindromic Repeats (CRISPR)/CRISPR-associated protein (Cas) system has been successfully repurposed for efficient genome editing in mammalian cells (Cong et al., 2013; Mali et al., 2013). This has opened the door to direct gene correction therapy for many inherited retinal diseases. Nevertheless, one of the greatest challenges is the efficient delivery of the CRISPR/Cas genome-editing system to the target tissues or cells in living organisms. Due to the large size of the commonly used SpCas9 (*Streptococcus pyogenes*, ∼4.2 kb) and the loading capacity of some currently available viral vectors for ocular gene therapy such as adeno-associated virus (AAV), recent studies have demonstrated that a dual AAV2 system can be used to deliver CRISPR/Cas9 to effectively perform DNA editing in retinal cells in adult mice (Bakondi et al., 2016; Hung et al., 2016; Latella et al., 2016; Ruan et al., 2017; Yu et al., 2017; Li et al., 2019). Despite the success of this dual-vector strategy, it is challenging to transduce two AAVs into one cell and clearly activity of the CRISPR/Cas9 machinery requires the receipt of both the endonuclease and sgRNA expression cassettes.

With the expanding repertoire of CRISPR/Cas endonucleases, various CRISPR/Cas systems have been developed that utilise smaller Cas endonuclease from different bacterial species, such as Cas12a (*Acidaminococcus*, ∼3.9 kb or *Lachnospiraceae*, ∼3.7 kb), SaCas9 (*Staphylococcus aureus*, 3.2 kb), CjCas9 (*Campylobacter jejuni*, 2.9 kb), NmCas9 (*Neisseria meningitidis*, ∼3.2 kb), making it possible to use a single vector to package both the Cas endonuclease and its sgRNA. A handful of studies have reported the successful *in vivo* genome editing of SaCas9(Maeder et al., 2019) CjCas9 (Kim et al., 2017; Jo et al., 2019), Cas12a (Koo et al., 2018), and NmeCas9 (Xia et al., 2018), in retinal cells. These various CRISPR/Cas systems differ in their editing efficacy, packagability and protospacer-adjacent motif (PAM) requirement (listed in Table S2), which largely expands the *in vivo* application of CRISPR/Cas based genome editing in various tissues or cells. There have been a small number of studies, which have applied all-in-one AAV vector mediated CRISPR/Cas genome editing in different cells including retinal pigment epithelium cells. Eunji and colleagues reported the successful disruption of the *Vegfa* or *Hif1a* genes in mouse RPE cells using single AAV-CjCas9 (Kim et al., 2017). Other groups have utilised a single AAV vector to deliver SaCas9, or NmeCas9(Ibraheim et al., 2018) to a variety of somatic tissue in mice (Ran et al., 2015; Ibraheim et al., 2018; Jarrett et al., 2018; Pan et al., 2018; Xu et al., 2019). Despite the encouraging *in vivo* application of these CRISPR/Cas systems, delivered via dual or all-in-one vectors, it is not clear which are the most efficacious for retinal editing *in vivo*.

The aim of this study was to directly compare the CRISPR/Cas endonuclease activity of single/dual AAV strategies for retinal cell gene editing in transgenic mice expressing a cre-sensitive yellow fluorescent protein (YFP) reporter. To achieve this, we designed YFP-targeting sgRNAs for each Cas endonuclease and quantified the editing efficiency, indicated by the disruption of YFP *in vitro* and *in vivo*.

## MATERIALS AND METHODS

### Ethics approval and animal maintenance

All experimental studies were performed in accordance with the Association for Research in Vision and Ophthalmology Statement for the Use of Animals in Ophthalmic and Vision Research and the requirements of the National Health and Medical Research Council of Australia (Australian Code of Practice for the Care and Use of Animals for Scientific Purposes). This study was approved by the Animal Ethics Committees of the University of Tasmania (Reference Number A0014827). CMV-Cre and Rosa26-YFP transgenic mouse lines were maintained on a C57BL/6 background and intercrossed to generate experimental offspring that were heterozygous for each transgene. At postnatal day (P) 0-2, offspring were genotyped using a BlueStar flashlight (Nightsea, Lexington, USA) to detect YFP expression in the brain, paws and ears. Adult (8-12 weeks old) CMV-Cre::Rosa26-YFP transgenic mice (YFP mice), which express YFP throughout the retina, were maintained and bred at University of Tasmania (Hobart, Australia). Animals were group housed with same-sex littermates in Optimice micro-isolator cages (Animal Care Systems, Colorado, USA) with uninhibited access to food and water. They were maintained on a 12 hour light (50 lux illumination) and 12 hour dark (<10 lux illumination) cycle, at 20°C to minimize possible light-induced damage to the eye.

### Design and construction of Cas endonucleases and sgRNAs vectors

Single guide RNAs targeting the same 5′ region of the *YFP* gene were designed using a CRISPR design tool (http://crispr.mit.edu) with different relevant PAM sites (Figure 1a). Briefly, three sgRNAs for SpCas9 (referred as SpCas9-YFP sgRNA1, 2, and 3), two sgRNAs with different lengths for Cas12a (referred as Cas12a-YFP sgRNA 20nt and 23nt), two sgRNAs for CjCas9 (referred CjCas9-YFP sgRNA1 and 2) and one sgRNA for SaCas9 (referred as SaCas9-YFP sgRNA, as only one possible PAM site was found in that region) were designed. These sgRNAs were then cloned into the pX552-CMV-mCherry vector, which was generated by replacing the hSyn1 promoter in the pX552-mCherry vector (Addgene No. 87916) with the CMV promoter. A control sgRNA, targeting the *LacZ* gene (5’-TGCGAATACGCCCACGCGAT-3’), was designed based on a previous study by Swiech et al (Swiech et al., 2015), and LacZ sgRNA plasmids were generated and used for *in vitro* validation.

**Figure 1.**
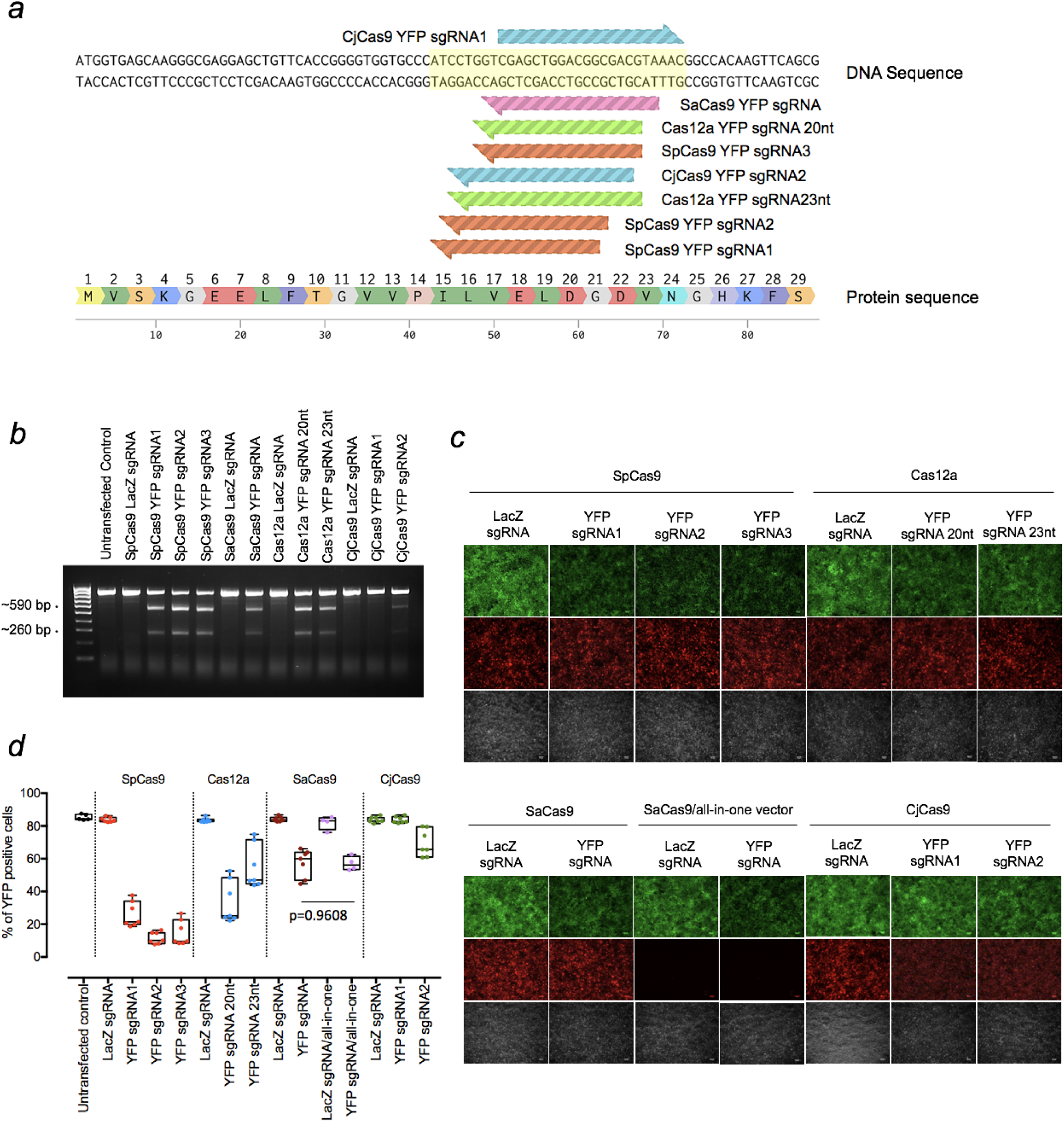
*In vitro* YFP sgRNA validation and selection. (a) *YFP*-targeting sequence for sgRNA design. *YFP*-targeting sgRNAs were designed (3 sgRNAs for SpCas9, 1 sgRNA for SaCas9, 2 sgRNAs for Cas12a and 2 for CjCas9). (b) T7E1 assay to detect cleavage efficiency for *YFP*. Expected cleavage products by T7E1 were detected in 2% TAE gel. * cleavage products around 590bp and 260bp. (c) Representative fluorescence microscopy images showing YFP expression in cells transfected with different CRISPR/Cas constructs. *Scale bar:100 µm.* (d) Flow cytometry analysis for sgRNA selection. Data are represented as mean ± SEM for 4-7 independent replicates. Intergroup comparisons were performed using a one-way ANOVA, and corrected for multiple comparisons. HEK293A cells without YFP expression were also included as negative control. No significant difference in *YFP* editing was observed between single and dual CRISPR/SaCas9 vector systems (p=0.9608). Selected sgRNAs for *in vivo* testing were SpCas9 YFPsgRNA2, Cas12a YFP sgRNA20nt and CjCas9 YFPsgRNA2.

Cas endonuclease plasmids were generated following different cloning approaches. The miniCMV-SpCas9 (SpCas9) construct was generated by replacing the CMV promoter with a miniCMV promoter in the pX551-CMV-SpCas9 plasmid (Addgene No. 107024) via AgeI and XbaI restriction enzyme sites. Other CRISPR/Cas endonucleases (SaCas9, Cas12a and CjCas9) were subcloned from pX601-SaCas9 (kindly provided by Feng Zhang; Addgene No. 61591), pcDNA3.1-hAsCpf1 (kindly provided by Feng Zhang; Addgene No. 69982) and pX404-CjCas9 (kindly provided by Feng Zhang; Addgene No. 68338) into pX551-CMV-SpCas9 plasmid by replacing SpCas9.

All-in-one single vector miniCMV-SaCas9/YFP sgRNA or miniCMV-SaCas9/LacZ sgRNA were generated based on the pX601-SaCas9 (kindly provided by Feng Zhang; Addgene No. 61591) by replacing the CMV promoter with miniCMV promoter and adding SpA terminator and inserting YFP sgRNA or LacZ sgRNA.

### Cell culture and transfection

HEK293A cells that stably express YFP (HEK293A-YFP) were generated as previously described (Hung et al., 2016, 2018). Cells were maintained in Dulbecco’s modified Eagle’s media (DMEM) (catalog no. 11965118; Life Technologies Australia, Mulgrave, VIC, Australia) supplemented with 10% fetal bovine serum (Sigma-Aldrich, St. Louis, MO, USA), 2 mM glutamine (catalog no. 2503008; Life Technologies Australia), antibiotic-antimycotic (catalog no. 15240062; Life Technologies Australia) in a humidified 5% CO_2_ atmosphere at 37°C. HEK293A-YFP cells were transfected with 750 ng of Cas endonuclease plasmid (under CMV promoter) and 750 ng of related sgRNA plasmid, or 750 ng of single SaCas9 plasmid, using lipofectamine 2000 (catalog no. 11668019; Life Technologies Australia), according to manufacturer’s instructions. YFP expression was evaluated 10 days later by collecting images of the HEK cell cultures using a fluorescent microscope and by performing a flow cytometric analysis. Genomic DNA was extracted from cells after each treatment and used to carry out a T7 endonuclease 1 (T7E1) assay (see below). The detailed information of reagents is provided in Table S1.

For Cas endonuclease detection, HEK293A cells were transfected with 1000 ng of the Cas endonuclease plasmid (under the miniCMV promoter) or the CjCas9 plasmid (pX551-CMV-CjCas9), and protein lysates were generated 2 days later to perform a Western blot analysis.

### Genomic DNA extraction and T7E1 mismatch detection assay

Genomic DNA was extracted with QuickExtract DNA Extraction Solution (catalog no. QE09050; Lucigen, Biosearch technologies, Middleton, WI, USA) and used as the DNA template for PCR reactions performed using KAPA HiFi HotStart DNA Polymerase (catalog no. KR0369; Roche Diagnostics Australia, North Ryde NSW, Australia) with primers listed in Table S3 (CMV SeqFWD forward and EYFP SURVEYOR reverse primers). PCR products were then denatured at 95°C for 10 min and gradually lowered to room temperature to allow for DNA heteroduplex formation, which were then digested by T7 Endonuclease I (catalog no. M0302S; New England Biolabs, Ipswich, MA, USA) following the manufacturer’s instructions. The digested products were visualized on 2% (w/v) agarose gels.

### Western blot analysis

To validate that Cas protein expression was being driven effectively by the Cas endonuclease plasmids, HEK293A cells were transfected with miniCMV-SpCas9, miniCMV-SaCas9, miniCMV-Cas12a, miniCMV-CjCas9 plasmids, and CMV-CjCas9 (under CMV promoter). Cells were collected at day 2 post transfection, and protein was extracted and quantified as described previously(Li et al., 2019). Protein samples were separated by using NuPAGE Electrophoresis system (Life Technologies Australia), after which proteins were transferred to polyvinylidene fluoride (PVDF) membranes (catalog no. 162-0177; Bio-Rad Laboratories; Hercules, CA, USA). Membranes were blocked with 5% (w/v) skim milk in TBS-T (10 mM Tris, 150 mM NaCl, and 0.05% Tween-20) at room temperature for 1 hour and then incubated with a mouse monoclonal HA antibody (F-7) (1:500 dilution; catalog no. sc-7392; Santa Cruz Biotechnology, Dallas, TX, USA) or mouse monoclonal β-actin antibody (1:1000 dilution; catalog no. catalog no. MAB 1501; Merck Millipore, Burlington, MA, USA) at room temperature for 1 hour. Membranes were washed, further incubated with a horseradish peroxidase-conjugated goat anti-mouse secondary antibody (1:5000 dilution; catalog no. A-11045; Life Technologies Australia) at room temperature for 1 h, and developed using the Amersham ECL Prime Western Blotting Detection Kit (catalog no. RPN2232; GE Healthcare Australia, Parramatta, NSW, Australia).

### Viral production

The AAV7m8 vectors were prepared by transfecting HEK293D cells (kindly provided by Ian Alexander, Children’s Medical Research Institute, Australia) with the miniCMV-Cas (SpCas9, SaCas9, Cas12a and CjCas9), CMV-CjCas9 or CMV-mCherry, selected YFP targeting sgRNAs or miniCMV-SaCas9/YFP sgRNA (all-in-one single vector) plasmids, helper plasmid (pXX6; kindly provided by Richard Samulski, The University of North Carolina School of Medicine, USA) and AAV2-7m8 capsid plasmid (7m8; Addgene No. 64839) using the calcium phosphate method (Hung et al., 2016). Viral vectors were purified using an AAVpro® Purification Kit (All Serotypes) (catalog no. 6666; Clontech Laboratories, Mountain View, CA, USA) 48 hours after viral transduction. Viral titrations were determined by real-time quantitative PCR using a Fast SYBR Green Master Mix (catalog no. 4385612; Life Technologies Australia) with AAV-ITR primers (Table S3). The titrations of AAV7m8 were provided in Table S4.

### Intravitreal injection

Mice were anesthetized with an intraperitoneal injection of ketamine (60 mg/kg) and xylazine (10 mg/kg). Bioccular, intravitreal injections were performed under a surgical microscope, using a hand-pulled glass needle connected to a 10 μL Hamilton syringe (Bio-Strategy, Broadmeadows, VIC, Australia), as described previously (Hung et al., 2016, 2018). Eyes with severe surgical or post-operative complications such as ocular hemorrhage or inflammation were excluded from the study. A scleral incision was made with a 30G needle before the glass needle was inserted into the vitreous to inject 1μL of the dual vector system (2.5×10^9^vg AAV7m8-Cas endonuclease and 2.5×10^9^vg AAV7m8-YFP sgRNA), the SaCas9 single vector system (2.5×10^9^vg AAV7m8-miniCMV-SaCas9/YFP sgRNA and 2.5×10^9^vg AAV7m8-mCherry) or the control vector (2.5×10^9^vg AAV7m8-mCherry). A total of 150 YFP transgenic mice were randomly allocated to the following groups: mCherry control (n=11), miniCMV-SpCas9 (n=12), miniCMV-SaCas9 (n=20), miniCMV-Cas12a (n=15), CMV-CjCas9 (n=11), miniCMV-CjCas9 (n=9) and miniCMV-SaCas9/YFP sgRNA (n=20), receiving the same viral injection regimen in each eye.

### Retinal flat mounts and histology

Enucleated eyes were immersion fixed in ice-cold 4% (w/v) paraformaldehyde in PBS for 1 hour before the retina was removed using a dissecting microscope as described previously(Hung et al., 2018). Processed retinal flat mounts were stained with NucBlue™ Live ReadyProbes™ Reagent (catalog no. R37605; Life Technologies Australia) for 20 minutes at room temperature before mounting with Dako Fluorescent mounting medium (catalog no. s3020; DAKO, Carpinteria, CA, USA). For histological assessment, eyes were fixed in 4% paraformaldehyde (w/v) in PBS for 1 hour and embedded in optimal cutting temperature compound (Leica Biosystems, Germany) and stored at -80C until cryosectioning. Serial 10 to 20-µm-thick cryosections were collected directly onto FLEX glass slides, followed by staining and mounting. Images of the retina were collected using an Olympus VS120 Slide Scanner or Perkin Elmer Spinning Disk Confocal Microscope (Zeiss spinning disk, Germany).

### Retinal dissociation and flow cytometry analysis

Retinas were rapidly dissected and digested using a papain dissociation kit (catalog no. LK003176; Worthington Biochemical Corporation, Lakewood, NJ) following the manufacturer’s instructions to obtain a homogenous cell suspension. After dissociation, retinal cells were resuspended in FACS buffer (1% Bovine Serum Albumin in Phosphate Buffered Saline) and stained with DAPI (5 µg/mL; catalog no. D1306; Life Technologies Australia) to exclude dead cells. Dissociated retinal cells from C57BL/6 mice were used as a negative control for YFP expression. Live retinal cells with mCherry (532nm, 622/22nm) and/or YFP (488nm, 513/26nm) expression were detected by flow cytometry (MoFlo ASTRIOS; Beckman Coulter, Brea, CA, USA). We quantified the proportion of mCherry-labelled cells that co-labelled for YFP in each retina using FlowJo analysis software (FlowJo®; FlowJo LLC, Ashland, OR, USA). Eyes with severe surgical complications such as cataract or retinal detachment or those with negligible mCherry expression were excluded from the final FACS analysis.

### Statistical Analysis

GraphPad Prism7 software (GraphPad Software, Inc., La Jolla, CA, USA) was used for statistical analyses. Data were represented as mean ± SEM and were analyzed using unpaired one-way analyses of variance (ANOVA). A value of p < 0.05 was considered statistically significant.

## RESULTS

### *In vitro* YFP sgRNA selection and Cas endonuclease validation

To select the most effective sgRNA for each Cas endonuclease, we first validated the on-target editing efficacy of different Cas endonucleases together with their respective sgRNAs using a T7E1 assay in HEK293A-YFP cells. Robust cleavage activity was evident in the groups transfected with the Cas endonuclease and their respective *YFP*-targetting sgRNAs, except for those treated with either CjCas9-*YFP* targeting constructs or *LacZ*-targeting controls (Figure 1b). Here, SpCas9-*YFP* targeting constructs were the most efficacious at knocking out *YFP* transgene expression, followed by Cas12a-*YFP* and SaCas9-*YFP* targeting constructs (Figure 1c).

The *YFP* disruption efficacy for each CRISPR/Cas construct was further quantified through flow cytometric analysis (Figure 1d). Compared to their non-targeting LacZ sgRNA counterparts, the percentage of *YFP*-expressing cells was significantly reduced by those transfected with SpCas9 and a *YFP*-targeting sgRNA (*YFP* sgRNA1: 26.0±2.9%, n=7, p<0.0001; *YFP* sgRNA2: 11.5±1.3%, n=7, p<0.0001; and *YFP* sgRNA3: 14.7±2.9%, n=7, p<0.0001). Similarly, Cas12a-targeting conditions resulted in appreciable *YFP* transgene knockout with a preference for a 20 nt-protospacer (*YFP* sgRNA 20nt: 33.6±4.9%, n=7, p<0.0001; and 23nt sgRNA: 55.0±5.0%, n=7, p<0.0001). Comparatively, CjCas9 was less effective at abrogating *YFP* transgene expression (*YFP* sgRNA2: 69.5±3.1%, n=7, p=0.0011) and failed to induce significant gene knockout in one of the conditions (CjCas9-*YFP* sgRNA1: 83.7±0.7%, n=7, p=0.999); while there was no significant difference in editing efficiency (p=0.9608) between the use of the SaCas9 single CRISPR construct (SaCas9/*YFP*-targeting sgRNA: 57.3±3.2%, n=4) and the dual CRISPR/Cas construct system (SaCas9 and its *YFP*-targeting sgRNA: 57.0±2.0%, n=7). The most effective *YFP*-targeting sgRNA for each Cas endonuclease (*YFP* sgRNA2 for SpCas9, 20nt *YFP* sgRNA for Cas12a and *YFP* sgRNA2 for CjCas9) were selected for subsequent *in vivo* testing.

To validate the protein expression of HA-tagged Cas endonuclease in a recombinant AAV vector (driven by minimal promoter, miniCMV or full-length CMV promoter), HEK293A cells were transfected with miniCMV-SpCas9, miniCMV-SaCas9, miniCMV-Cas12a, miniCMV-CjCas9 and CMV-CjCas9. Cas endonuclease protein expression was evident with the use of the minimal promoter, except for CjCas9 (miniCMV-CjCas9; Figure S1), which required the full-length CMV promoter to drive transgene expression (CMV-CjCas9). Therefore, four miniCMV-Cas endonuclease (SpCas9, SaCas9, and Cas12a) constructs and the CMV-CjCas9 construct were used along with their selected sgRNAs for further *in vivo* CRISPR/Cas editing comparison.

### *In vivo* AAV7m8 delivery of CRISPR/Cas in the mouse retina

AAV7m8-mediated gene expression (mCherry) and distribution were assessed on retinal sectioning/flatmounts of the CMV-Cre::*Rosa26*-*YFP* mouse eye 5 months after intravitreal injection (Figure 2a). Retinal flatmount images from AAV7m8-CRISPR/Cas-injected retina showed a wide distribution of mCherry expression (Figure 2b). Fluorescence images revealed AAV7m8 transduction (as indicated by mCherry expression) was visible throughout the retina, including the Ganglion Cell Layer (GCL), Inner Nuclear Layer (INL) and even some parts of the retinal outer nuclear layer (ONL), with major expression within INL (Figure 2c). Moreover, *YFP* expression could be found in all the layers of the retina with no observable difference between AAV7m8-CRISPR/Cas-treated mice and control mice.

**Figure 2.**
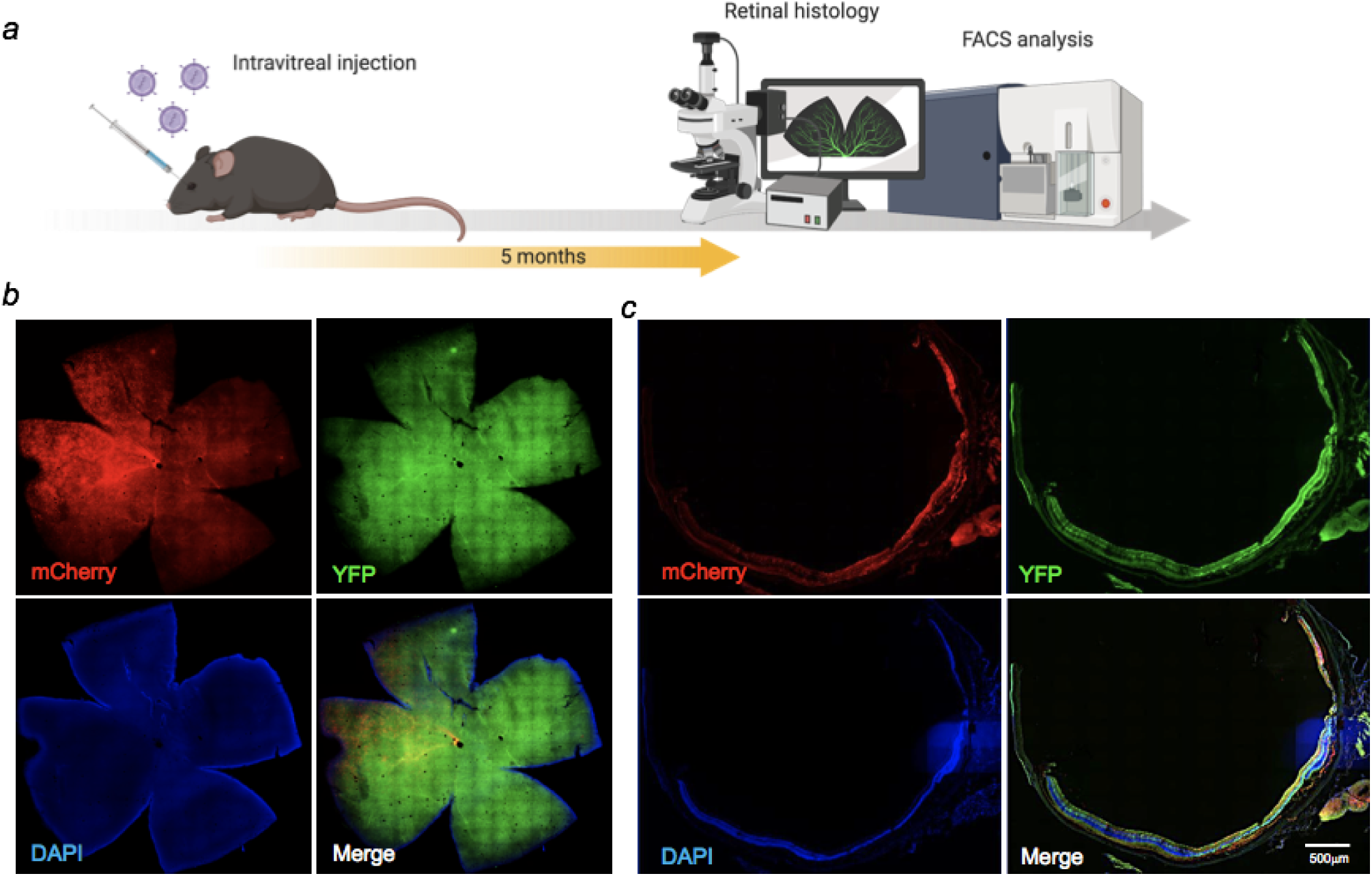
AAV7m8 mediated delivery of CRISPR/Cas to the mouse retina *in vivo*. (a) Schematic diagram of *in vivo* experiment. Mice were sacrificed 5 months after intravitreal injection. (b) Representative cross section image from retina co-transduced with AAV7m8-miniCMV-CjCas9 and its selected YFP sgRNA. Robust AAV7m8 transductions in the retina were found. Scale bar: 200 µm. Images were taken by a Zeiss spinning disk confocal microscope. (c) Representative retinal whole-mount images from two mouse eyes receiving different AAV7m8-CRISPR/Cas. Left panel: AAV7m9-CMV-CjCas9, mouse ID 29, right eye; Right panel: AAV7m8-miniCMV-CjCas9, mouse ID 76, right eye. Scale bar: 500 µm. Images were taken by Olympus Slide Scanner.

### Comparison of *YFP* knockout in the mouse retina with different Cas endonucleases Constructs

Five dual CRISPR/Cas constructs (miniCMV-SpCas9, miniCMV-SaCas9, miniCMV-Cas12a, miniCMV-CjCas9, and CMV-CjCas9) with their selected *YFP*-targeting sgRNA and a single all-in-one SaCas9 CRISPR construct (miniCMV-SaCas9/YFP-targeting sgRNA) were used to compare the editing efficacy in retinal cell *in vivo* (Figure 3a). To evaluate and compare the *YFP* knockout *in vivo* delivered by AAV7m8-mediated different CRISPR/Cas system, the percentage of *YFP* disruption among mCherry positive retinal cells was quantified by flow cytometry (Figure 3b). The flow cytometric gating strategy is shown in Supplementary data (Figure S2). Representative dot plots in Figure 3b illustrate the difference in *YFP* disruption in retinal cells receiving AAV7m8-SpCas9 CRISPR vector or control vector. Differences in AAV7m8 transduction efficiency were observed between CRISPR/Cas treatment groups (Figure 3c) with a lower percentage of mCherry positive cells observed in the retinas transfected with AAV7m8-Cas12a (35.1±2.8%, n=15) and CjCas9 (28.6±3.4%, n=9) CRISPR vectors. Retinas receiving AAV7m8-SpCas9 and SaCas9 (both single and dual vector system) CRISPR vectors had a relatively high proportion of mCherry expression (50.0±4.6%, n=12; 52.0±4.3%, n=20; 57.7±3.3%, n=19 respectively). For *YFP* disruption, AAV7m8-SpCas9 CRISPR vector (18.9±2.9%, n=12) had the highest knockout efficiency of YFP among all the CRISPR/Cas systems, followed by SaCas9 (single vector system: 8.4±3.4%, n=20; dual vector system: 9.8±2.6%, n=20) and Cas12a (5.4±2.0%, n=15), while CjCas9 showed no disruption of *YFP* expression (Figure 3d). Moreover, there was no significant difference in the *YFP* disruption in the retinas receiving single and dual SaCas9 CRISPR constructs (single vector system: 8.4±3.4% vs dual vector system: 9.8±2.6%, n=20, p=0.9994) (Figure 3d). Despite efficient in vivo YFP knockout in animals administered SpCas9, SaCas9 and Cas12a constructs, there was a high degree of variability between individual animals within identical treatment groups.

**Figure 3.**
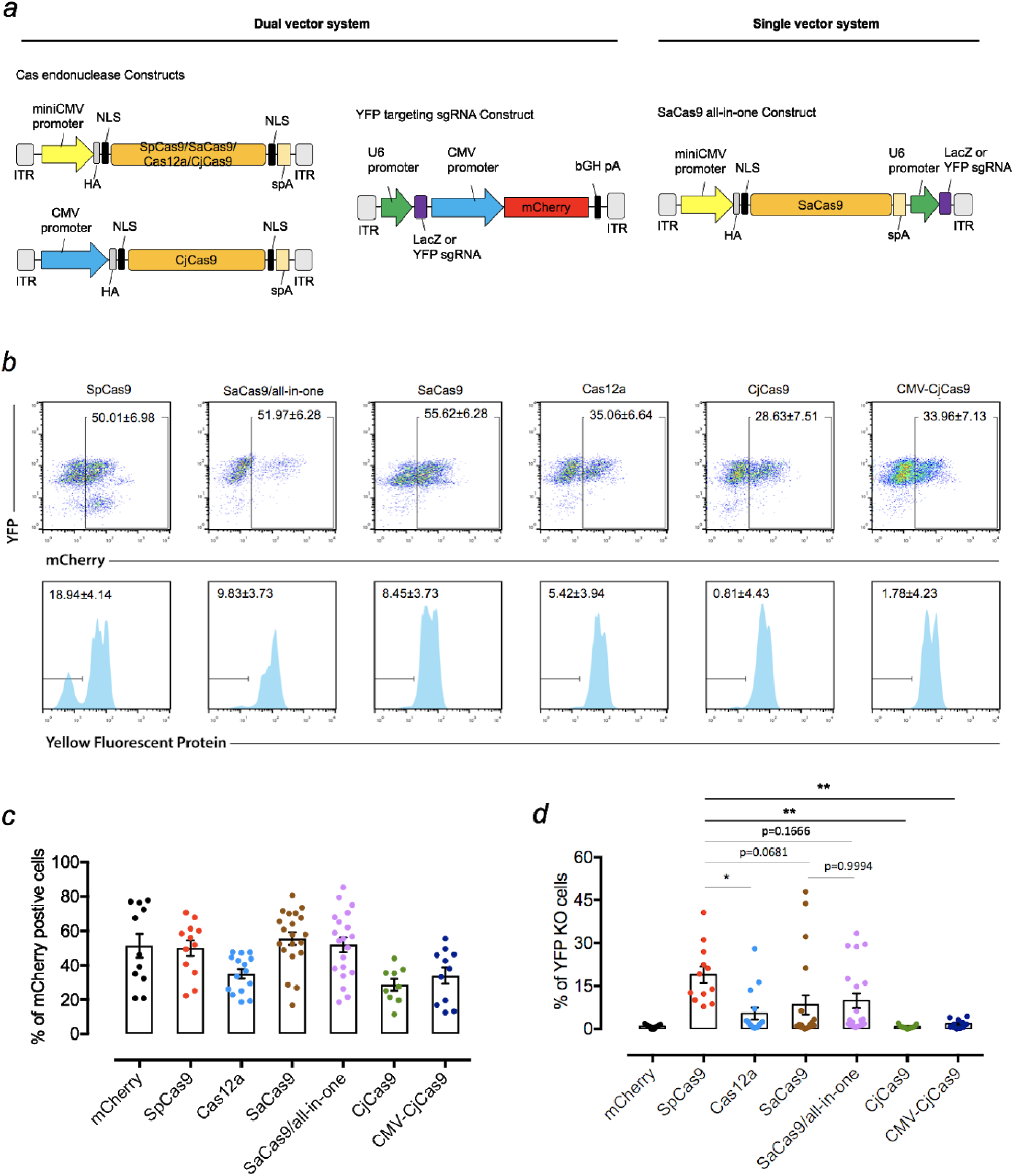
Comparison of YFP disruption in retinal cells with different CRISPR/Cas systems delivered by AAV7m8. (a) Schematic of the dual and single vector systems. For dual vector plasmids, the Cas endonuclease was driven by miniCMV or CMV promoter (only miniCMV promoter driven plasmids were packaged into AAV for *in vivo* testing), whilst the sgRNA was driven by U6 promoter and mCherry under the control of CMV promoter to confirm vector transfection. For the single vector systems, an all-in-one plasmid with SaCas9 or CjCas9 was designed with the Cas endonuclease being driven by a miniCMV promoter and sgRNA by U6 promoter. For Cas12a, we used the Cas endonuclease from *Acidaminococcus* (originally designated AsCpf1). A hemagglutinin (HA) tag was fused to the C-terminus of Cas endonuclease in the vector. (b) Representative FACS plots of dissociated retinal cells receiving different AAV7m8. AAV7m8-mCherry control (left panel) or AAV7m8-SpCas9 / AAV7m8-YFP sgRNA (right panel). The histograms in the lower panels (lower panel) were based on mCherry gating. Dissociated cells from one retina were used in each group. (c) Comparison of AAV7m8 transduction in the retina indicated by mCherry expression by FACS. mCh: mCherry, SP: SpCas9, SSA: Single SaCas9, DSA: Dual SaCas9, CMV-CJ: CMV-CjCas9, CJ: CjCas9. Data are presented as mean ± SEM for 9-20 independent samples in each group. Statistical analysis between groups was performed using one-way ANOVA followed by multiple comparisons test. n=number of injected eyes. * *p*<0.05, ** *p*<0.01, *** *p*<0.001. (d) Comparison of *YFP* disruption in mCherry positive cells by FACS. Data are presented as mean ± SEM for 9-20 independent samples in each group. Non-parametric one-way ANOVA – the Kruskal-Wallis test was applied as data in two groups (Cas12a and Dual SaCas9) did not pass the D’Agostino & Pearson normality test. n=number of injected eyes. * *p*<0.05, ** *p*<0.01, *** *p*<0.001.

## DISCUSSION

In this study, we provide a direct comparison of the efficacy for retinal editing *in vivo* with four different currently available CRISPR/Cas systems. Here, we showed that SpCas9 and Cas12a achieved better knockout efficiency than SaCas9 and CjCas9 *in vitro*. AAV7m8-packaged CRISPR/Cas construct with SpCas9 was found to have the highest editing efficacy among all Cas endonucleases *in vivo*. No significant difference in *YFP* gene editing was found between single and dual CRISPR/SaCas9 vector systems *in vitro* and *in vivo*.

This study was based on our previous work, which used AAV2-mediated delivery of CRISPR/Cas9 to achieve efficient gene editing in the inner layer of retina in Thy1-*YFP* mice (Hung et al., 2016). To assess and compare the genome efficiency in the whole retina, we applied a different murine model CMV-Cre::*Rosa26*-*YFP* transgenic mice (*YFP* mouse) which express YFP throughout the retina. To this end, we used the AAV7m8-pseudotype, an AAV2-based variant with enhanced retinal transduction when delivered through intravitreal injection (Dalkara et al., 2013; Khabou et al., 2016). As the degeneration of RPE and photoreceptors are involved in the majority of inherited retinal diseases, efficient gene delivery of CRISPR constructs to the outer layer of retina is imperative for therapeutic retinal gene editing. Subretinal injection of conventional AAVs (e.g. AAV2) has high photoreceptor transduction rate, but it is surgically challenging with more complications. In addition, the cellular transduction is confined within the injection bubble of the retina. Our study shows that AAV7m8-mediated CRISPR/Cas has reasonable pan-retinal transduction.

The stringent design of this study ensured a fair comparison of editing efficiency between different CRISPR/Cas systems. First, we analysed the *YFP* coding sequence for all potential PAM sites for each Cas endonuclease and then designed sgRNA targeting *YFP* within a similar region. We additionally employed the same ubiquitous promoter (CMV for *in vitro* sgRNA selection, miniCMV for *in vivo*) for each endonuclease, and employed for the same virus. The only exception to this design was the use of the more potent CMV promoter for *in vivo* CjCas9 constructs, due to its poor expression on western blots of *in vitro* HEK293A cells. Despite this modification, CjCas9 barely demonstrated *YFP* knockout on flow cytometric analysis of *in vivo* specimens. We hypothesize that variation in CjCas9 codon-optimization may account for the differences observed in study compared to that reported by other groups (Kim et al., 2017).

We additionally found differences in gene knockout efficiency between *in vitro* and *in vivo* modes. For the *in vitro* study, SpCas9 outperformed Cas12a, followed by SaCas9 and CjCas9. For *in vivo* samples, SpCas9 remained the best-performing Cas endonuclease among all, without a clear trend among the other Cas orthologs. Initially, we hypothesized that the single all-in-one SaCas9 vector expressing both the SaCas endonuclease and its respective sgRNA may have a competitive or even higher editing efficiency compared to dual-vector mediated-editing with SpCas9, but we did not observe this result in our *in vivo* test.

In summary, we demonstrate that AAV7m8-mediated delivery of CRISPR/SpCas9 construct achieves the most efficient gene modification in retinal cells *in vitro* and *in vivo* among four currently available CRISPR/Cas systems.

## ACKNOWLEDGEMENTS

This work was supported by funding from a Bayer Global Ophthalmology Award, the Ophthalmic Research Institute of Australia, an Australian National Health and Medical Research Council (NHMRC) grant (APP1123329), an NHMRC Practitioner Fellowship (AWH, 1103329), a NHMRC Career Development Award (KMY, APP1045240) and an NHMRC Research Fellowship NHMRC Senior Research Fellowship (AP, 1154389). Centre for Eye Research Australia receives Operational Infrastructure Support from the Victorian Government.

## AUTHOR CONTRIBUTIONS

FL, AWH, GS and SSCH conceptualised and designed this study. FL, KW, JHW, CDL, JB, JC, QW, QL, and PNT performed all laboratory-based experiments. KY, RCBW, AP and ALC provided reagents and resources for this work. FL wrote the original draft, with all authors providing reviewing and editing. GS and AWH jointly supervised this work.

## SUPPLEMENTARY MATERIALS

**Table S1.** Comparison of Cas orthologs for *in vivo* retinal gene editing application.

**Table S2.** Sequence of primers for sgRNA cloning, vector construction, sequencing and qPCR analysis.

**Table S3.** AAV7m8 titrations.

**Figure S1**. The *in vitro* validation of Cas endonuclease expression.

**Figure S2.** Representative FACS plot showing gating strategy.

**Table S1.** Summary of reagents and resources used.

| REAGENT or RESOURCE | SOURCE | IDENTIFIER |
| --- | --- | --- |
| **Antibodies** |  |  |
| HA-probe Antibody (F-7) | Santa Cruz Biotechnology | sc-7392 |
| mouse monoclonal β-actin antibody | Millipore | MAB 1501 |
| HRP-conjugated goat anti-mouse secondary antibody | Life Technologies Australia | A-11045 |
| **Chemicals** |  |  |
| Sodium azide ($\text{NaN}_3$) | Sigma-Aldrich | S2002-100g |
| Trizma base | Sigma-Aldrich | T1503-1KG |
| Glycine | Sigma-Aldrich | G8898-1KG |
| $\text{NaH}_2\text{PO}_4$ | Sigma-Aldrich | S0751-500G |
| Calcium chloride dihydrate ($\text{CaCl}_2 \cdot 2\text{H}_2\text{O}$) | Sigma-Aldrich | 223506-500G |
| Agarose | Sigma-Aldrich | A9539-250G |
| LB broth base | Life Technologies Australia | 12780-029 |
| LB broth with agar | Sigma-Aldrich | L2897-1KG |
| DAPI | Life Technologies Australia | D1306 |
| **Critical Commercial Assays** |  |  |
| TaKaRa AAVpro Purification Kit (All Serotypes) | Clontech Laboratories | 6666 |
| Fast SYBR Green Master Mix | Life Technologies Australia | 4385612 |
| Amersham ECL Prime Western Blotting Detection Kit | GE Healthcare Australia | RPN2232 |
| QIAquick PCR purification kit (250) | Qiagen | 28106 |
| Qiaprep spin miniprep kit (250) | Qiagen | 27106 |
| HiSpeed Plasmid Maxi Kit (10) | Qiagen | 12262 |
| QIAfilter Plasmid Mega Kit (5) | Qiagen | 12281 |
| **Experimental Models: Cell Lines** |  |  |
| Human embryonic kidney (HEK) 293A | Life Technologies Australia | R70507 |
| HEK293A-YFP | Guei-Sheung Liu Lab | n/a |
| HEK293D | Ian Alexander Lab | n/a |
| **Experimental Models: Organisms/Strains** |  |  |
| CMV-Cre::Rosa26-YFP transgenic mice | University of Tasmania | n/a |
| **Plasmids** |  |  |
| pX552-mCherry | Feng Zhang Lab | Addgene No. 87916 |
| pX601-AAV-CMV::NLS-SaCas9-NLS-3xHA-bGHpA;U6::BsaI-sgRNA | Feng Zhang Lab | Addgene No. 61591 |
| pY010 (pcDNA3.1-hAsCpf1) | Feng Zhang Lab | Addgene No. 69982 |
| pX404-CjCas9 | Feng Zhang Lab | Addgene No. 68338 |
| pX551-CMV-SpCas9 | Alex Hewitt Lab | Addgene No. 107024 |
| pXX6 | Richard Samulski Lab | n/a |
| 7m8 | John Flannery & David Schaffer Lab | Addgene No. 64839 |

| Softwares and Algorithms |  |  |
| --- | --- | --- |
| ImageJ version 1.48 | Schneider et al., 2012 | https://imagej.nih.gov/ij |
| GraphPad Prism7 | GraphPad Software | http://graphpad.com |
| Adobe Photoshop (CC 2017.1.1) | Adobe | https://adobe.com |
| Guide RNA design | Feng Zhang Lab | http://crispr.mit.edu |
| Guide RNA design | Benchling | https://benchling.com |
| FlowJo | FlowJo LLC | https://flowjo.com |
| Others |  |  |
| Dulbecco's modified Eagle's media (DMEM) | Life Technologies Australia | 11965118 |
| Iscove's Modified Dulbecco's Medium (IMDM) | Life Technologies Australia | 12440061 |
| Glutamine | Life Technologies Australia | 2503008 |
| Antibiotic-antimycotic | Life Technologies Australia | 15240062 |
| Opti-MEM | Life Technologies Australia | 11058021 |
| Lipofectamine 2000 transfection reagent | Life Technologies Australia | 11668019 |
| Virkon (5gm tablet) | MedShop Australia | MED1610661 |
| AgeI | New England Biolabs | R0552L |
| XbaI | New England Biolabs | R0145L |
| T7 Endonuclease I (T7E1) | New England Biolabs | M0302S |
| QuickExtract DNA Extraction Solution | Lucigen | QE09050 |
| KAPA HiFi HotStart DNA Polymerase | Roche Diagnostics<br>Australia | KR0369 |
| Polyvinylidene fluoride (PVDF) membranes | Bio-Rad Laboratories | 162-0177 |
| TE buffer (PH 8.0, 500 mL) | Life Technologies<br>Australia | AM9849 |
| Sodium Dodecyl Sulfate (SDS) | Life Technologies<br>Australia | 15525017 |
| Tween 20 | Sigma-Aldrich | P9416-50ML |
| SYBR Safe-DNA Gel Stain | Life Technologies<br>Australia | S33102 |
| 1KB plus DNA ladder | Life Technologies<br>Australia | 10787018 |
| Gel loading Dye Purple (6x) | New England Biolabs | B7025S |
| NuPAGE® LDS Sample Buffer (4X) | Life Technologies<br>Australia | 0007 |
| Novex® sharp pre-stained protein standard | Life Technologies<br>Australia | LC5800 |
| Cell Lysis Buffer | Thermo Fisher Scientific | 89900 |
| T4 DNA ligase | New England Biolabs | M0202S |

**Table S2.**
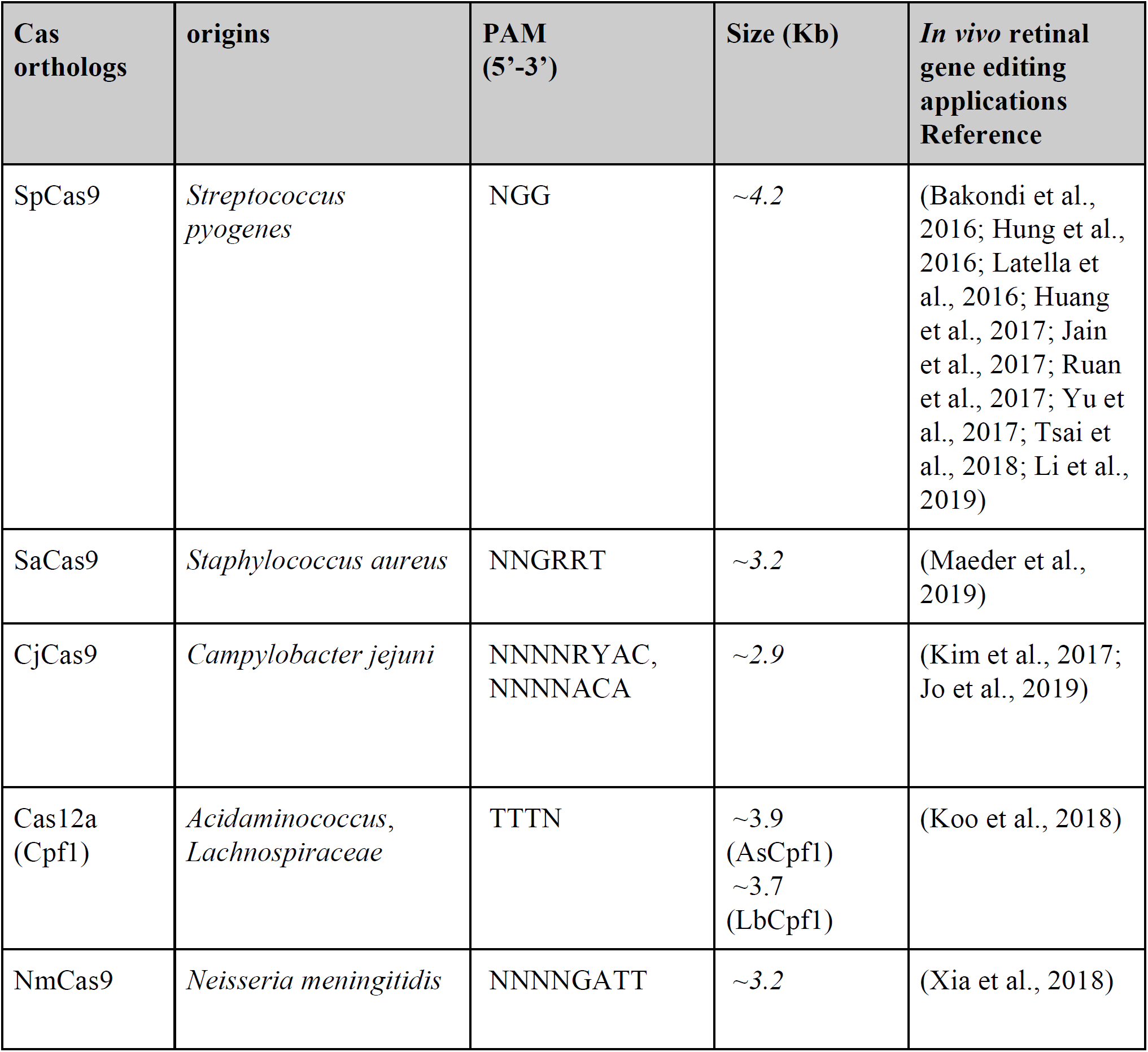
Comparison of main Cas orthologs for *in vivo* retinal gene editing applications.

| Cas orthologs | origins | PAM (5'-3') | Size (Kb) | *In vivo* retinal gene editing applications<br>Reference |
| --- | --- | --- | --- | --- |
| SpCas9 | *Streptococcus pyogenes* | NGG | ~4.2 | (Bakondi et al., 2016; Hung et al., 2016; Latella et al., 2016; Huang et al., 2017; Jain et al., 2017; Ruan et al., 2017; Yu et al., 2017; Tsai et al., 2018; Li et al., 2019) |
| SaCas9 | *Staphylococcus aureus* | NNGRRT | ~3.2 | (Maeder et al., 2019) |
| CjCas9 | *Campylobacter jejuni* | NNNNRYAC,<br>NNNNACA | ~2.9 | (Kim et al., 2017; Jo et al., 2019) |
| Cas12a (Cpf1) | *Acidaminococcus*,<br>*Lachnospiraceae* | TTTN | ~3.9 (AsCpf1)<br>~3.7 (LbCpf1) | (Koo et al., 2018) |
| NmCas9 | *Neisseria meningitidis* | NNNNGATT | ~3.2 | (Xia et al., 2018) |

**Table S3.**
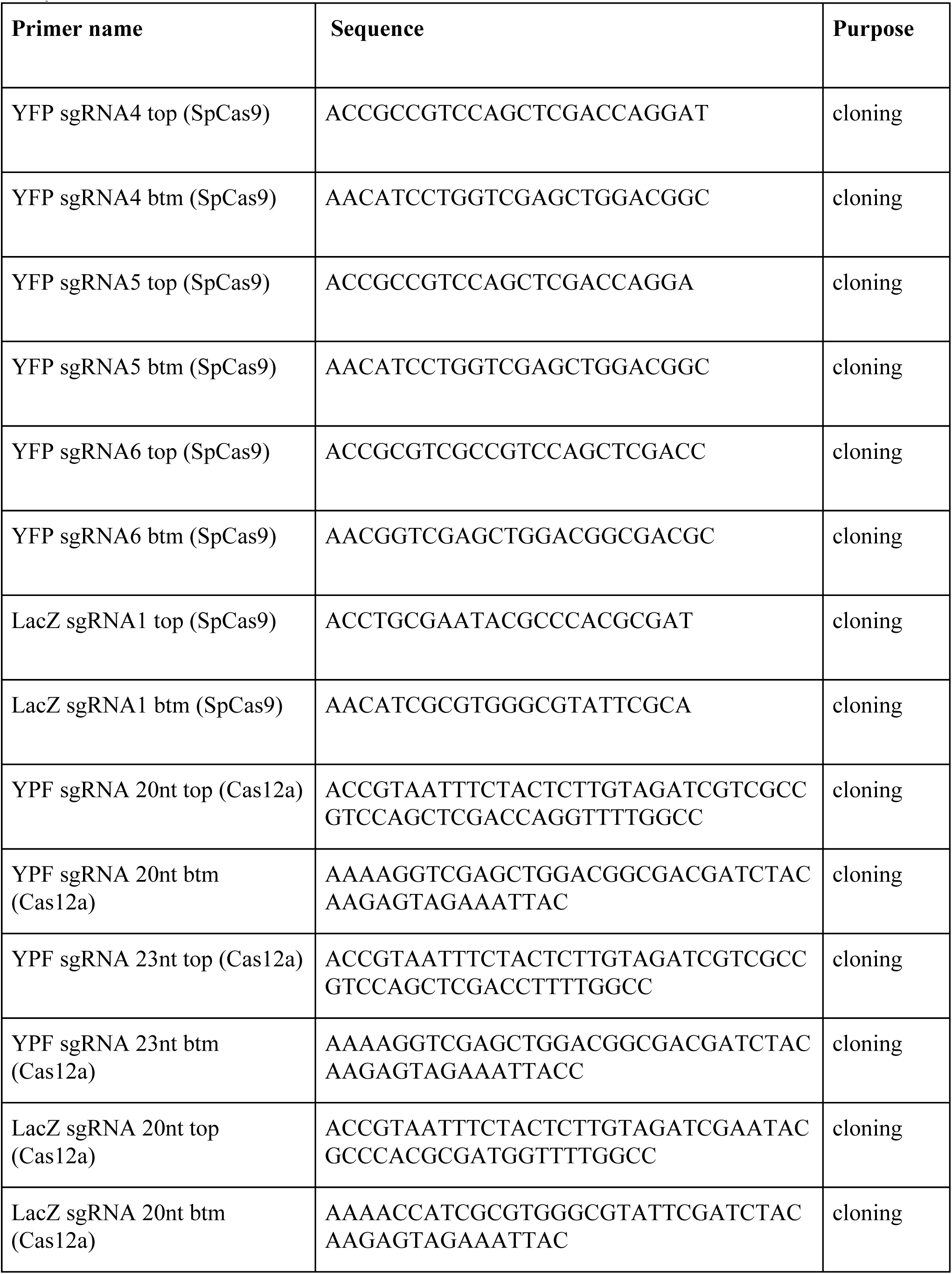
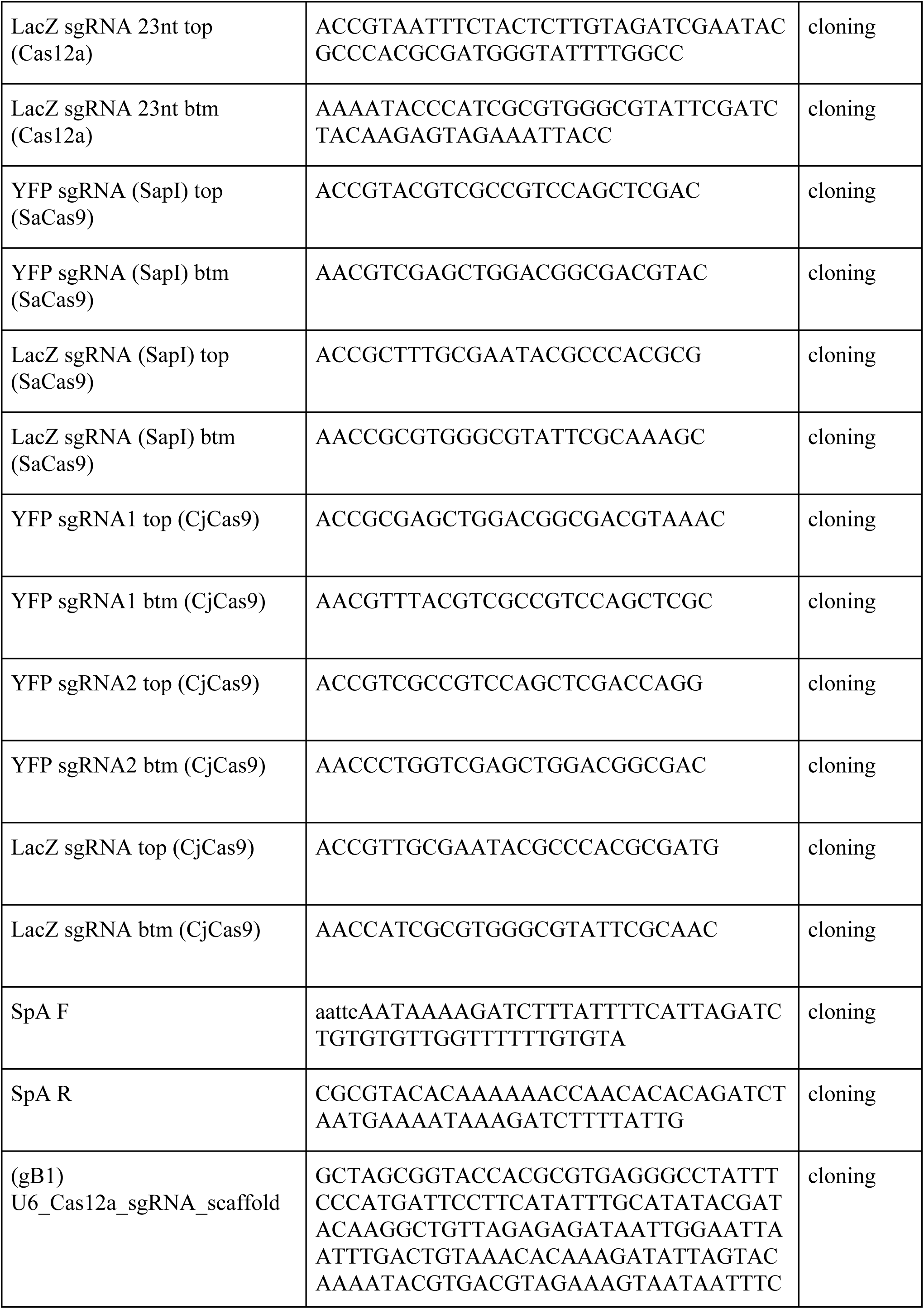
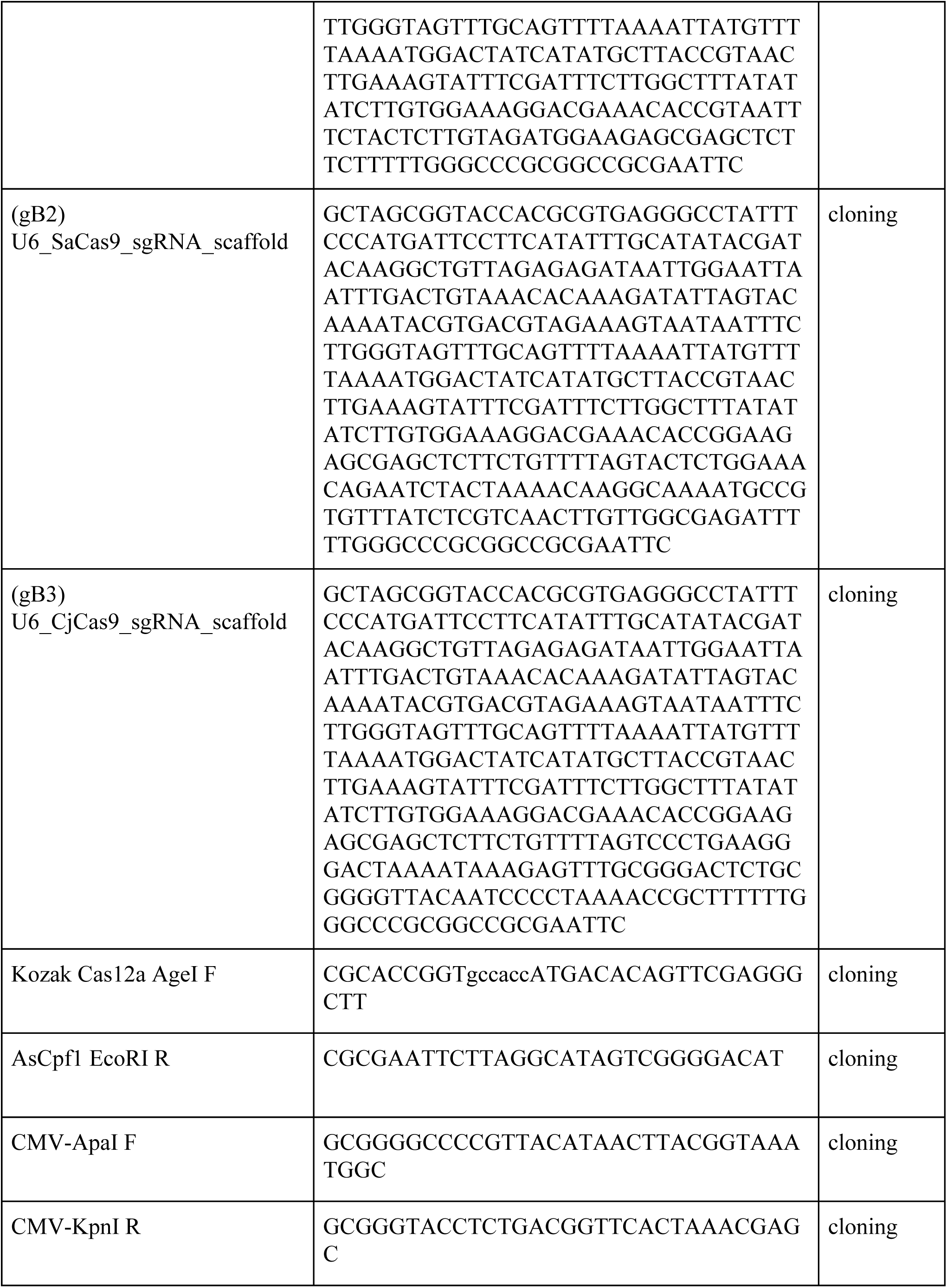
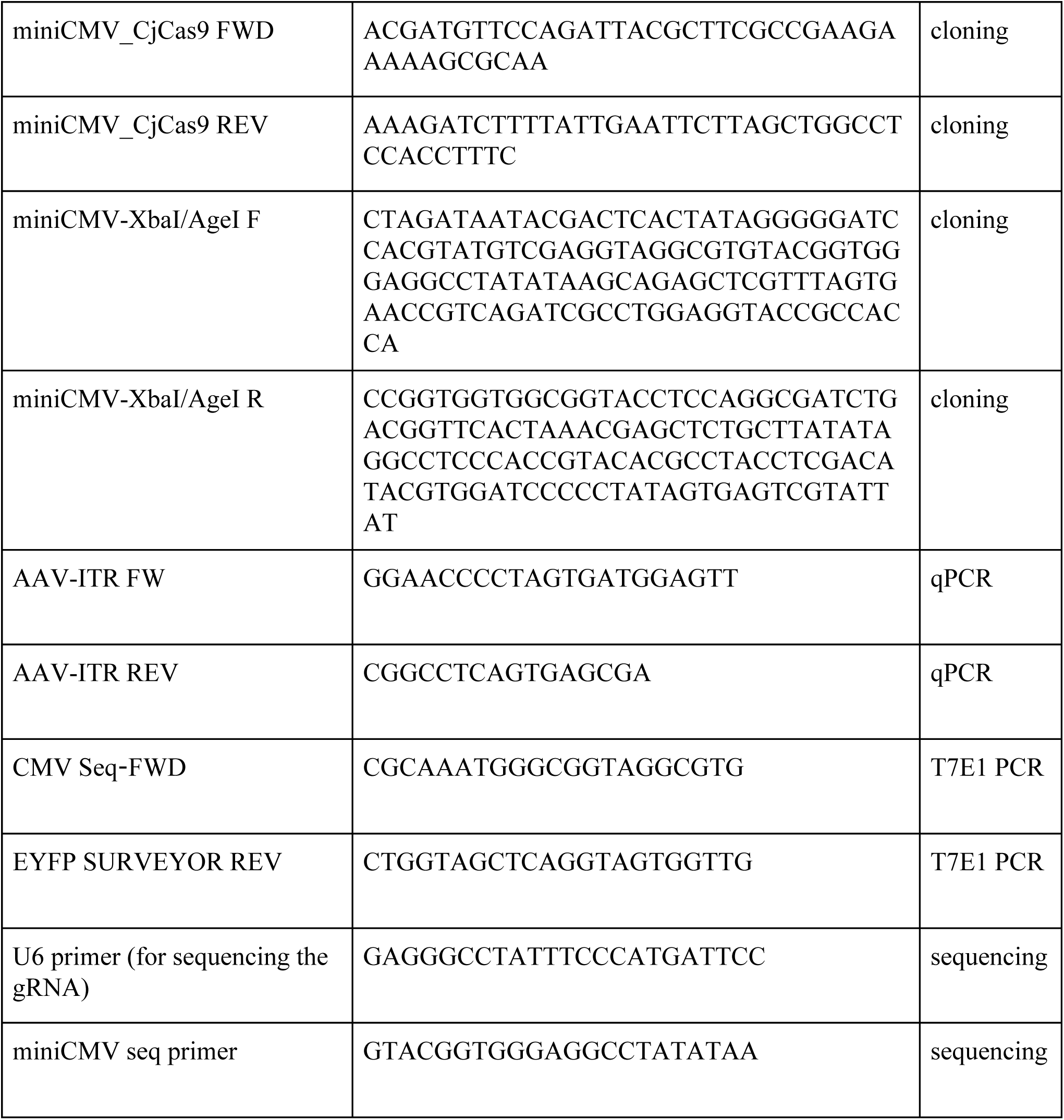
Sequence of primers for sgRNA cloning, vector construction, sequencing and qPCR analysis.

**Table S4.** AAV7m8 titrations.

| **AAV7m8** | **Vector** | **Titration (vg/mL)** |
| --- | --- | --- |
| AAV7m8-SpCas9 | pX551-miniCMV-SpCas9 | $5.13 \times 10^{14}$ |
| AAV7m8-SaCas9 | pX551-miniCMV-SaCas9 | $5.48 \times 10^{14}$ |
| AAV7m8-Cas12a | pX551-miniCMV-Cas12a | $6.22 \times 10^{14}$ |
| AAV7m8-CjCas9<br>(CMV) | pX551-CMV-CjCas9 | $1.61 \times 10^{15}$ |
| AAV7m8-CjCas9<br>(miniCMV) | pX551-miniCMV-CjCas9 | $3.53 \times 10^{14}$ |
| AAV7m8-YFP<br>sgRNA2 (SpCas9) | pX552-CMV-mCherry-YFP sgRNA2<br>(SpCas9) | $2.31 \times 10^{14}$ |
| AAV7m8-YFP sgRNA<br>20nt (Cas12a) | pX552-CMV-mCherry-YFP sgRNA 20nt<br>(Cas12a) | $1.06 \times 10^{15}$ |
| AAV7m8-YFP sgRNA<br>(SaCas9 dual) | pX552-CMV-mCherry-YFP sgRNA<br>(SaCas9) | $1.22 \times 10^{14}$ |
| AAV7m8-YFP<br>sgRNA2 (CjCas9) | pX552-CMV-mCherry-YFP sgRNA2<br>(CjCas9) | $1.26 \times 10^{15}$ |
| AAV7m8-YFP sgRNA<br>(SaCas9 single) | pX601-miniCMV-SaCas9-U6-YFP sgRNA | $2.55 \times 10^{14}$ |
| AAV7m8-mCherry | pX552-CMV-mCherry | $1.58 \times 10^{14}$ |

**Figure S1.**
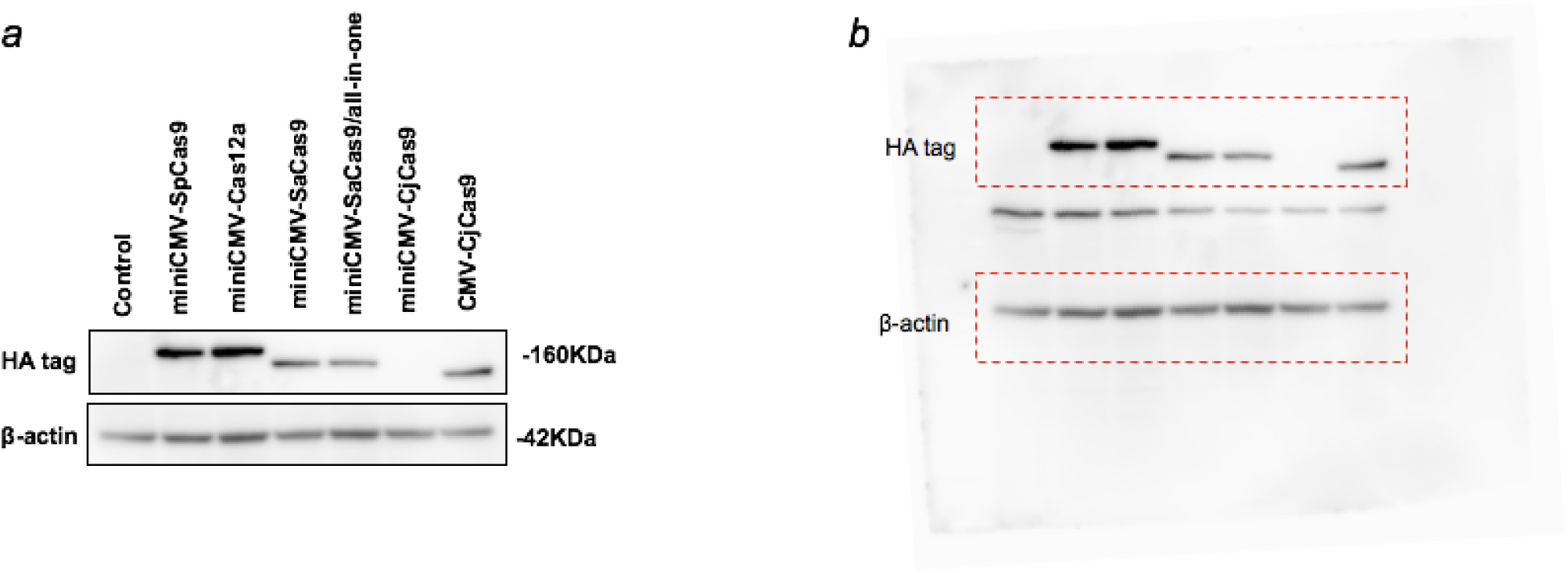
Cas endonuclease validation. *In vitro* validation of different Cas endonucleases by western blot. Representative western blot of Cas protein expression in HEK293A cells treated with AAV-Cas plasmids two days after transfection. CjCas9 expression was not detectable in cells transfected with miniCMV-CjCas9 plasmid by western blot with HA tag antibody. β-actin was used as a loading control.

**Figure S2.**
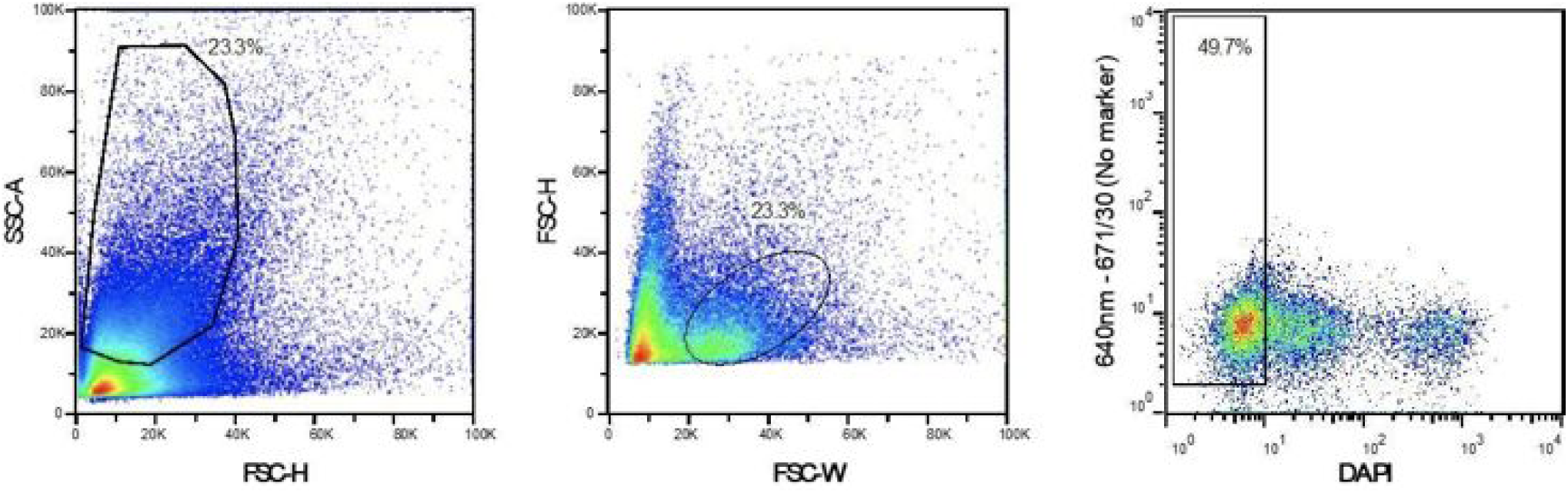
Representative FACS plot showing gating strategy. Single cells from dissociated retina were gated on forward scatter (FSC-H)/side scatter (SSC-A) plot and live cells were further gated on DAPI.

